# Rapid, but limited, zooplankton adaptation to simultaneous warming and acidification

**DOI:** 10.1101/2021.04.07.438881

**Authors:** Hans G. Dam, James A. deMayo, Gihong Park, Lydia Norton, Xuejia He, Michael B. Finiguerra, Hannes Baumann, Reid S. Brennan, Melissa H. Pespeni

## Abstract

Predicting the response of marine metazoans to climate change is hampered by a lack of studies on evolutionary adaptation, particularly to combined ocean warming and acidification (OWA). We provide evidence for rapid adaptation to OWA in the foundational copepod species, *Acartia tonsa*, by assessing changes in population fitness based on a comprehensive suite of life-history traits, using an orthogonal experimental design of nominal temperature (18°C, 22°C) and *p*CO_2_ (400, 2000 μatm) for 25 generations (~1 year). Egg production and hatching success initially decreased under OWA, resulting in a 56% reduction in fitness. However, both traits recovered by the third generation and average fitness was reduced thereafter by only 9%. Antagonistic interactions between warming and acidification in later generations decreased survival, thereby limiting full fitness recovery. Our results suggest such interactions constrain evolutionary rescue and add complexity to predictions of the responses of metazoan populations to climate change.

---

The continuous increase in atmospheric CO_2_ is unparalleled in the last 300 million years^1^, causing rapid ocean warming (OW)^2,3^ and ocean acidification (OA)^4^. Warming and acidification in coastal regions are projected to be more extreme^5,6^. Overall, novel environments may arise as a result of these changes^7^. Predicting how biota respond to this rapid global change is a crucial yet formidable scientific challenge, particularly whether populations can persist at similar levels and phenotypes in the future^8^. Organisms can respond to novel environments through phenotypic plasticity or evolutionary adaptation, both of which can mitigate the deleterious effects of climate change^8–12^. In particular, long-term experimental evolution studies are a powerful tool to examine the role of adaptation in mitigating the effects climate change on biota and to explore whether populations can evolve fast enough to keep pace^13^.

Organismal performance typically decreases sharply when temperatures increase beyond the optimum^14^ leading to disproportional deleterious effects on performance. In addition, marine metazoans require the mobilization of energy-demanding acid-base regulatory processes to counteract internal pH reduction to maintain homeostasis under high CO_2_ conditions. This may result in increased metabolic costs at the expense of growth and reproduction, even for non-calcifying metazoans^15–17^. Although factorial assessments of species sensitivities to warming and acidification have increased rapidly over the past years, multi-generational studies on metazoan populations responding to future simultaneous warming and acidification (OWA) are rare^18–20^. Moreover, lack of population fitness measures in existing studies preclude considerations of evolutionary rescue – evolution occurring sufficiently fast to allow population recovery before extirpation^21–23^.

We used an experimental evolution approach to test whether a marine zooplankter, the copepod *Acartia tonsa* (Dana; 1849), can adapt to environments created by OW, OA, and OWA conditions, to identify the functional traits under selection, and to assess evolutionary rescue. As the most abundant metazoans on the planet^24,25^, copepods link primary producers and other microbes to upper trophic levels, thereby influencing fisheries productivity^26,27^ and mediating marine biogeochemical cycles^28^. Specifically, *Acartia tonsa* is a dominant copepod in estuarine systems from tropical to temperate regions^29^ and a main prey item of forage fish^30^, which makes this species an important zooplankton model. Using both improvements in trait performance and population fitness across generations, we show rapid, yet limited copepod adaptation to combined ocean warming and acidification likely driven by an antagonistic interaction of warming and acidification.

## Experimental design

We measured five fitness-relevant life-history traits (survival, egg production rate (EPR), egg hatching success (HS), development time, and sex ratio) across 25 generations in an orthogonal design with two levels of CO_2_ and temperature. A population of *Acartia tonsa* was collected from Long Island Sound (41.3°N, 72.0°W; Groton, CT, USA) and kept under standard laboratory conditions (see methods) for at least three generations prior to the experiment. Four lines of the population were established with four replicates of each condition. The target (actual ± standard deviation) conditions were: ambient temperature = 18°C (18 ± 0.34, N = 330), ambient *p*CO_2_ = 400 μatm (379 ± 36, N = 18; pH = 8.26 ± 0.1, N = 330); high temperature = 22°C (22 ± 0.81, N = 336), and high *p*CO_2_ = 2000 μatm (2301 ± 215, N = 18; pH = 7.55 ± 0.08, N = 330). Ambient target levels represented extant conditions for this species in NE Atlantic estuaries (see methods for choice of temperature), and high levels corresponded to future conditions based on global projections for the years 2100-2300^1–4^, although *A. tonsa* already periodically experiences high temperature and CO_2_ levels in its growth season in NE Atlantic estuaries^31,32^. Summaries of the temperature, pH, and CO_2_ data are shown in Supplementary Tables 1-3. Details of statistical tests and their significance for the transgenerational experiment are in the methods section.

## Evidence for adaptation

During the first experimental generation (generation zero), EPR and HS declined in all three future (OW, OA, and OWA) conditions relative to ambient (AM) conditions: OW (EPR: *p* < 0.0001; HS: *p* < 0.01; t-test), OA (EPR: *p* < 0.0001; HS: *p* = 0.19; t-test), and OWA (EPR: < 0.0001; HS: *p* < 0.0001; t-test; Fig. 1), illustrating the ecological effects of these climate change variables. The decrease was strongest under OWA, particularly for HS, indicating synergistic deleterious effects of temperature and CO_2_.

**Figure 1.**
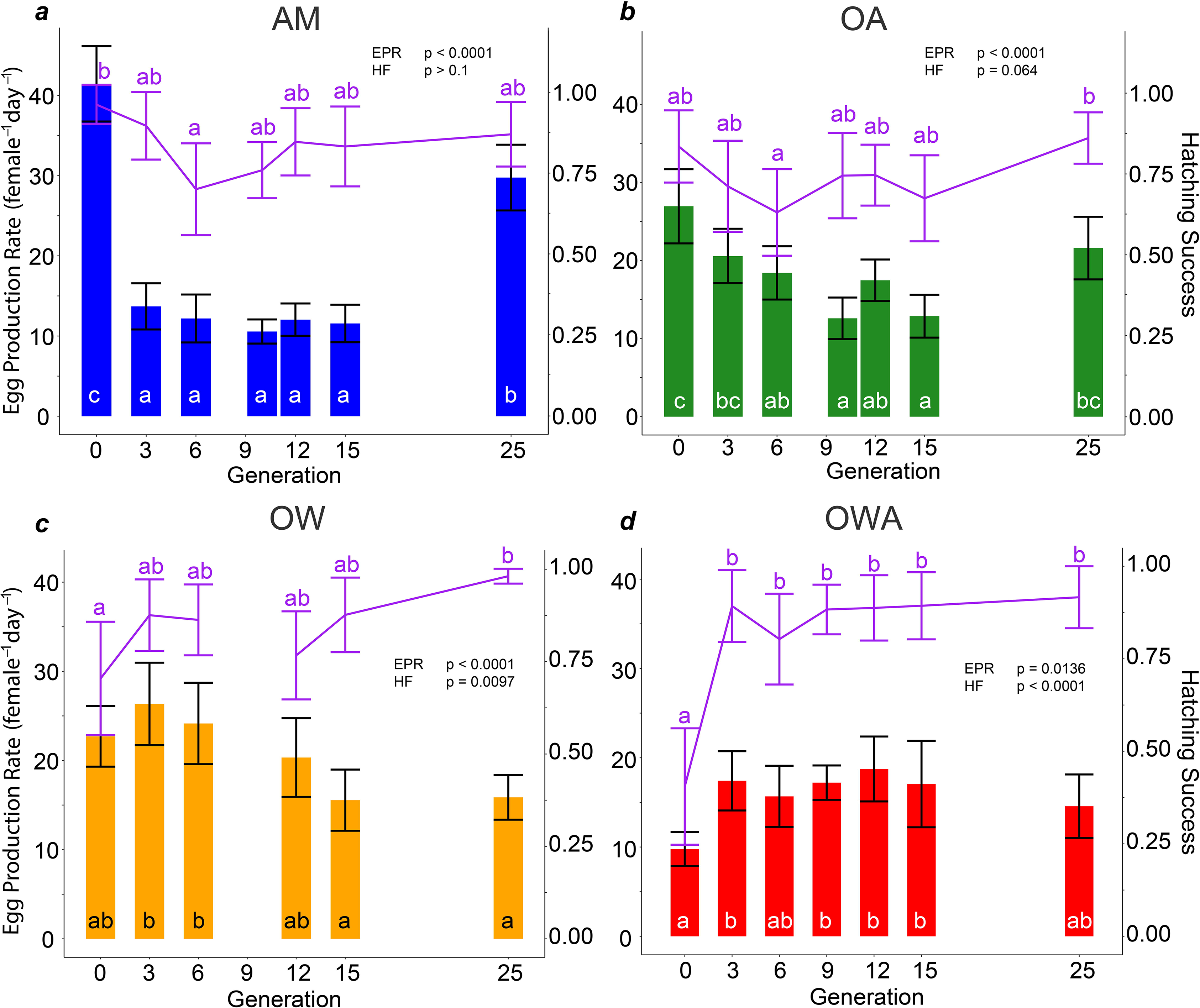
Mean egg production and mean hatching success for transgenerational experiment. Egg production rate (EPR, histograms, left Y axis) and egg hatching success (HS, lines, right Y axis) during the transgenerational experiment. **A**) Ambient conditions, AM: Blue (top left) – ambient temperature (18 °C) and CO_2_ (400 μatm); **B**) Ocean acidification conditions, OA: Green (top right) – ambient temperature and high CO_2_ (2000 μatm); **C**) Ocean warming conditions, OW: Orange (bottom left) – high temperature (22 °C) and ambient CO_2_; **D**) Ocean warming and acidification, OWA: Red (bottom right) – high temperature and high CO_2_. Error bars represent 95% confidence intervals. Probability (p) values for EPR and HS for each panel are derived from the effect of generation on each trait taken from generalized additive models. Letters represent statistically similar groupings for post-hoc t-test comparison. Box and whiskers plots of the same graphs are displayed on SI Fig. 6.

Significant interactions of temperature and CO_2_ on HS were evident across all generations (*p* < 0.0001; three-way ANOVA). After the first generation, EPR decreased under AM (*p* < 0.0001; GAM ANOVA) and OA conditions (*p* < 0.0001; GAM ANOVA), but partially recovered in later generations. Meanwhile, HS remained stable (AM GAM ANOVA: *p* = 0.14; OA GAM ANOVA: *p* = 0.064; Fig. 1). Under OW, EPR decreased with generation (*p* < 0.0001; GAM ANOVA) but did not recover. Meanwhile, HS increased for OW (*p* < 0.0001; GAM ANOVA; Fig. 1C) suggesting some degree of adaptation. By contrast, under OWA, EPR increased by 50% (*p* < 0.02; GAM ANOVA) and HS doubled (*p* < 0.0001; GAM ANOVA; Fig. 1D) by generation 3, and the improvements were maintained until generation 25. These changes yielded significant effects on population fitness (Table 1) and are consistent with rapid adaptation.

**Table 1.** Generational effects on fitness. Zero-inflated model results of generation and treatment on fitness, with replicates as random effects. Count model results represents when λ=0 values are omitted. Control treatment is the reference intercept for the model. Interactive effect of generation on AM conditions is noted as just “Generation” for each model results.

| *Predictors* | *Estimates* | *CI* | *P* |
| --- | --- | --- | --- |
| **Count model** |  |  |  |
| (Intercept) | 1.19 | 1.16-1.23 | **<0.001** |
| Generation | -0.00 | -0.00 – -0.00 | **0.001** |
| OA | 0.01 | -0.04 – 0.06 | 0.817 |
| OW | 0.04 | -0.01 – 0.09 | 0.115 |
| OWA | 0.13 | 0.07 – 0.18 | **<0.001** |
| Generation × OA | 0.00 | -0.00 – 0.00 | 0.951 |
| Generation × OW | 0.00 | 0.00 – 0.00 | **0.009** |
| Generation × OWA | -0.01 | -0.01 – -0.01 | **<0.001** |
| **Zero-inflated model** |  |  |  |
| (Intercept) | -1.97 | -2.62 – -1.32 | **<0.001** |
| Generation | -0.01 | -0.04 – 0.03 | 0.681 |
| OA | 0.55 | -0.36 – 1.46 | 0.233 |
| OW | 1.27 | 0.39 – 2.16 | **0.005** |
| OWA | 1.54 | 0.65 – 2.44 | **0.001** |
| Generation × OA | -0.03 | -0.07 – 0.02 | 0.212 |
| Generation × OW | -0.04 | -0.09 – -0.00 | **0.038** |
| Generation × OWA | -0.19 | -0.25 – -0.13 | **<0.001** |
| **Random effects** |  |  |  |
| $\sigma^2$ | 0.01 | | |
| $\tau_{00}$ Replicate | 0.00 | | |
| Intra-correlation coefficient | 0.08 |  |  |
| N Replicate | 16 |  |  |
| Observations | 2867 |  |  |
| Marginal $R^2$/Conditional $R^2$ | 0.230/0.293 | | |

Survival from freshly hatched nauplii to reproductively mature adults was independent of treatment (*p* > 0.05, two-way ANOVA)for the first 12 generations (Fig. 2). By generation 15, however, survival decreased by 30% under OW (*p* = 0.02, t-test) and OWA (*p* = 0.01, t-test) relative to ambient conditions. Acidification alone elicited no decrease in survival (*p* > 0.1, one-way ANOVA). Although survival recovered in the OW treatment by generation 25, survival decreased under OWA by an additional 50% relative to generation 15 for the same treatment (*p* < 0.001, t-test), demonstrating an antagonistic interaction between OW and OA in later generations^33^. Traits under selection are expected to increase towards phenotypic optima^34–36^, thus survival does not appear to be a trait under selection for adaptation to OW, OA or OWA conditions. High temperature resulted in faster development in both the OW and the OWA treatments (Supplementary Fig. 1) with 22-24% shorter development times than at ambient temperature (*p* < 0.0001; two-way ANOVA). High CO_2_, in contrast, resulted in 5-6% slower development times than ambient CO_2_ across all generations (*p* < 0.0001; two-way ANOVA). Thus, warming and acidification acted antagonistically on development time. Finally, a ~ 1:1 sex ratio remained unchanged for three of the four treatments (Supplementary Fig. 2). Under OWA, however, the proportion of females decreased across generations, with a significantly lower proportion in generation 25 than generation 0 (*p* < 0.01; Tukey HSD).

**Figure 2.**
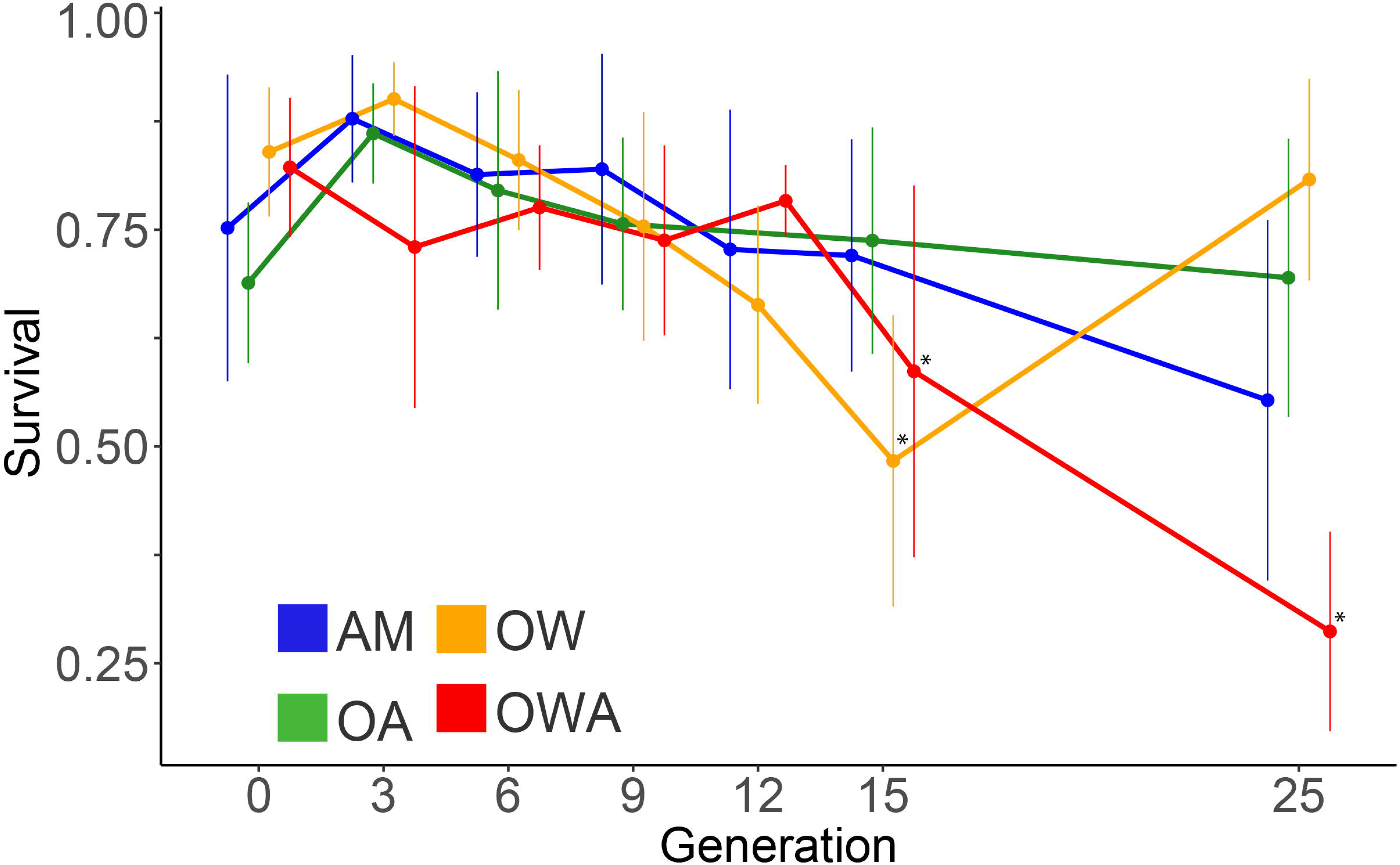
Survival for transgenerational study. Mean survival is for nauplius stage 1 to adult stage. Error bars represent 95% confidence intervals. Asterisks indicate significant decrease (*p* < 0.001). Treatment colors are the same as histograms in Fig. 1. Treatment curves are offset for clarity. Box and whiskers plots of the same graphs are displayed on SI Fig. 7.

To understand adaptation not just in terms of individual traits, but with respect to population fitness, we integrated all measured traits (survival, EPR, HS, development time, and sex ratio) to estimate the net reproductive rate, lambda^37^(λ), which is the fraction of the population replaced in a generation (Fig. 3). In generation 0, OWA conditions resulted in a 56% reduction in λ relative to ambient conditions, while OW and OA resulted in 23% and 13% reductions relative to ambient conditions, respectively (*p* < 0.01, t-test; Fig. 3). However, by generation 3, λ in OWA conditions had improved by 120% relative to generation 0 (*p* < 0.0001; Tukey HSD; Fig. 3) and recovered to levels equal to ambient conditions (*p* = 1.0; Tukey HSD; Fig. 3), mainly driven by improved hatching success (Fig. 1). The λ-frequency distribution shows an inflation of zero values (Supplementary Fig. 3) due to the high abundance of mate pairs with low HS at generation 0, particularly under OWA conditions. In later generations, as HS increased so did the probability of non-zero λ, and this was especially evident under OWA conditions (Supplementary Fig. 4). Thus, as generations progressed, selection under OWA conditions culled off low-fitness individuals in the population. Lambda remained high in OWA conditions until generation 15, after which there was a 19% reduction (*p* < 0.05, t-test; Fig. 3). This decrease was driven by reduced survival (Fig. 2), which suggests an inability of *Acartia tonsa* to maintain multiple optimal phenotypes, i.e., high hatching success and high survivorship, under OWA conditions. While λ was 22% lower under OWA than ambient conditions by generation 25, it was significantly higher than in generation 0 (*p* < 0.0001, t-test; Fig. 3), still consistent with adaptation over time. In accord, significant effects of generation on λ were evident under OW (*p* < 0.04; ANOVA) and OWA (*p* < 0.001; ANOVA) conditions, but not under ambient (*p* = 0.681; ANOVA) or OA (*p =* 0.212; ANOVA) conditions (Table 1). The latter suggests that OA alone was not a strong selective force in our study.

**Figure 3.**
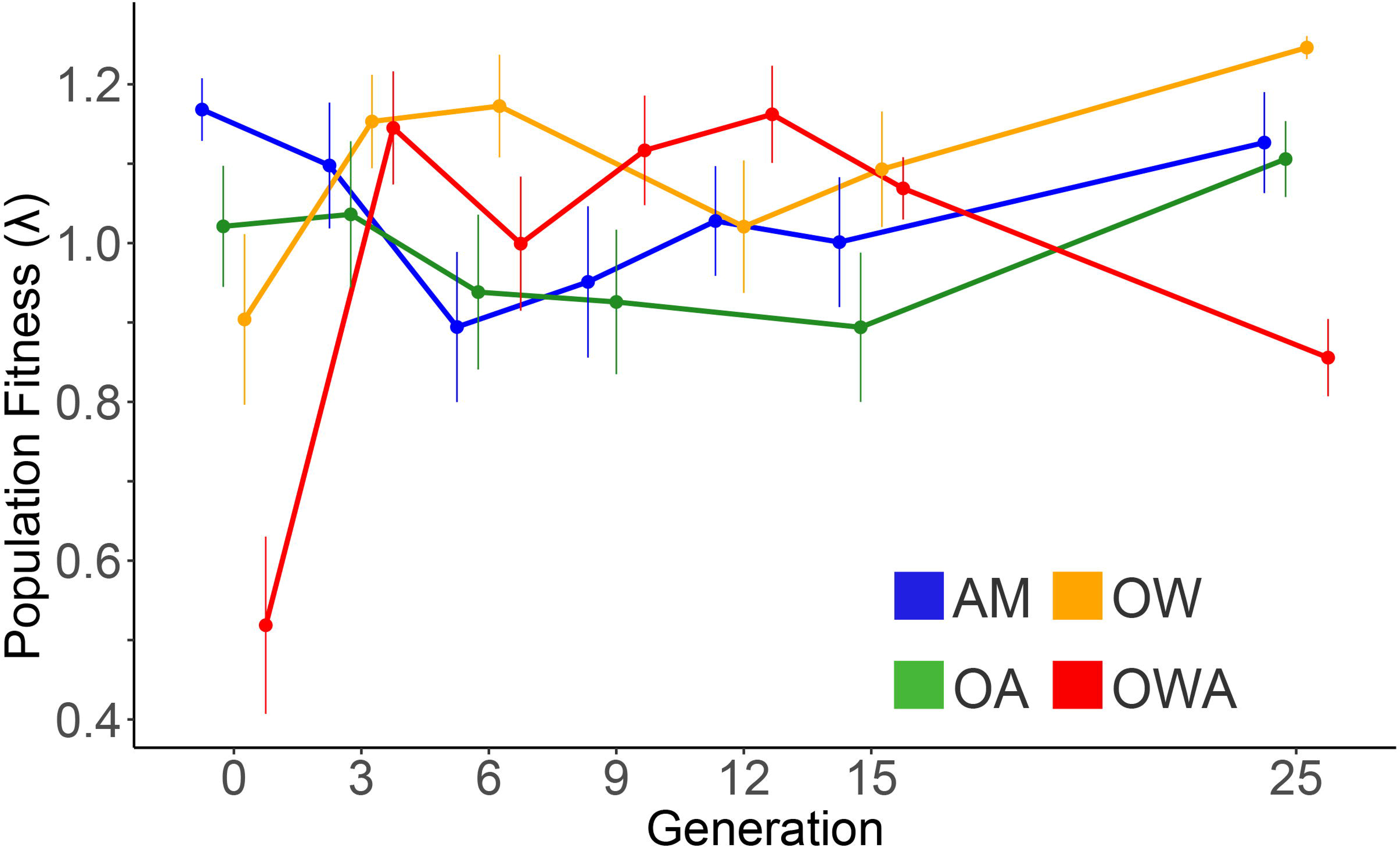
Fitness. Mean fitness values, λ, calculated for the transgenerational study. Error bars represent 95% confidence intervals. Treatment colors are the same as histograms in Fig. 1. Treatment curves are offset for clarity. Box and whiskers plots of the same graphs are displayed on SI Fig. 8.

Adaptation is evident when performance increases towards optimal phenotypes that increase fitness over time^34–36^. Here, we observed improved performance and fitness under OW and OWA conditions, but with important differences between the treatments: fitness was fully recovered under OW, but not under OWA conditions. *Acartia tonsa* exhibits strong tolerance to high temperatures, consistent with its seasonal dominance in the summer and wide-ranging latitudinal distribution^29,38^. However, the combination of elevated temperature and CO_2_ are known to affect ectotherm resource partitioning and energy distribution^15^, which might account for the limited recovery under OWA conditions. During adaptation, multiple phenotypes are expected to reach optimal levels of performance concurrently to yield the maximum possible population fitness for a particular environment. We hypothesize that under OWA conditions copepods could not sustain multiple optimized phenotypes, as evidenced by the observed reduction in survival following HS and EPR recovery to levels equal to or greater than those of ambient conditions. Under OW, copepods improved both EPR and HS (Fig. 1) and maintained high survival (Fig. 2) across generations to yield the highest λ level relative to all conditions by generation 25 (Fig. 3). By contrast, under OWA conditions the improvements in EPR and HS across generations yielded the highest λ values between generations 3 and 12 but decreases in survival reduced λ afterwards. These results are consistent with adaptation via selection, while simultaneously allowing maladaptive traits to persist^35^ under OWA conditions.

Genetic drift affects global, genome-wide patterns of genetic diversity, while selective processes affect specific regions of the genome^39,40^. We tested for signatures of genetic drift caused by potential bottlenecks by exploring patterns of nucleotide diversity (*π*) among treatments using single nucleotide polymorphisms identified with pooled sequencing of genomic DNA from each replicate of each treatment at generation 25. We found that global levels of nucleotide diversity were equivalent across treatment groups (Supplementary Figure 5; Wilcoxon Rank Sum test, *p* > 0.05; average *π* across treatments: AM: 0.0135 ± 0.006; OA: 0.0130 ± 0.006; OW: 0.0134 ± 0.006; OWA: 0.0138 ± 0.006), revealing no evidence of genetic bottlenecks or drift after 25 generations. In separate work, we observed divergence in allele frequencies and gene expression at generation 20 between the AM and OWA lines^41^. Among the four lineages at generation 25, allele frequency estimates indicate differentiation between the four lines despite the similar levels of nucleotide diversity (our own unpublished observations). Altogether, the improvement in traits and population fitness across generations coupled with the genetic differentiation between the OWA and the AM treatments are consistent with evolutionary adaptation in the OWA treatment.

## Traits affecting fitness

To assess the contribution of each life-history trait to adaptation, we quantified the strength of selection via the standardized linear selection coefficient of the fitness landscape^42^. High linear coefficients of selection for a given fitness landscape suggest that a particular trait is under selection, though not necessarily determining fitness. The coefficients are calculated from multiple regression models of relative λ against all changing life-history traits (see methods). We found that HS, but not EPR or survival, was under selection and had a positive impact on fitness. Specifically, relative λ increased as a function of HS with standardized linear selection coefficients (β) of 0.91 (AM), 0.94 (OA), 0.96 (OW), and 0.78 (OWA) at generation 0 (*p* < 0.0001, ANOVA; Fig. 4A). In addition, βHS was significantly different for all treatments between generations 0 and 25 (*p* < 0.0001, ANOVA), indicating changing degrees of selection on HS for all treatments between the first and last generations. By contrast, neither EPR or survival were significant at generation 0 (Fig. 4B, C; *p* > 0.1, ANOVA) suggesting that selection acted in favor of higher HS, but not towards higher survival or EPR for the OW and OWA treatments. By generation 25, the impact of HS on relative fitness decreased relative to generation 0 by 88% and 10% for the OW and OWA treatments, respectively. Such decreases corresponded to a shift in phenotype distribution towards higher HS and suggested that selection had relaxed by generation 25 as HS within the population reached the phenotypic optimum (Fig. 4A). Finally, despite the significant decrease in sex ratio over the 25 generations in the OWA treatment (Supplementary Fig. 2), sex ratio had no effect on relative fitness between generations 0 and 25 (*p* = 1.0, ANOVA), suggesting this trait was not under selection.

**Figure 4.**
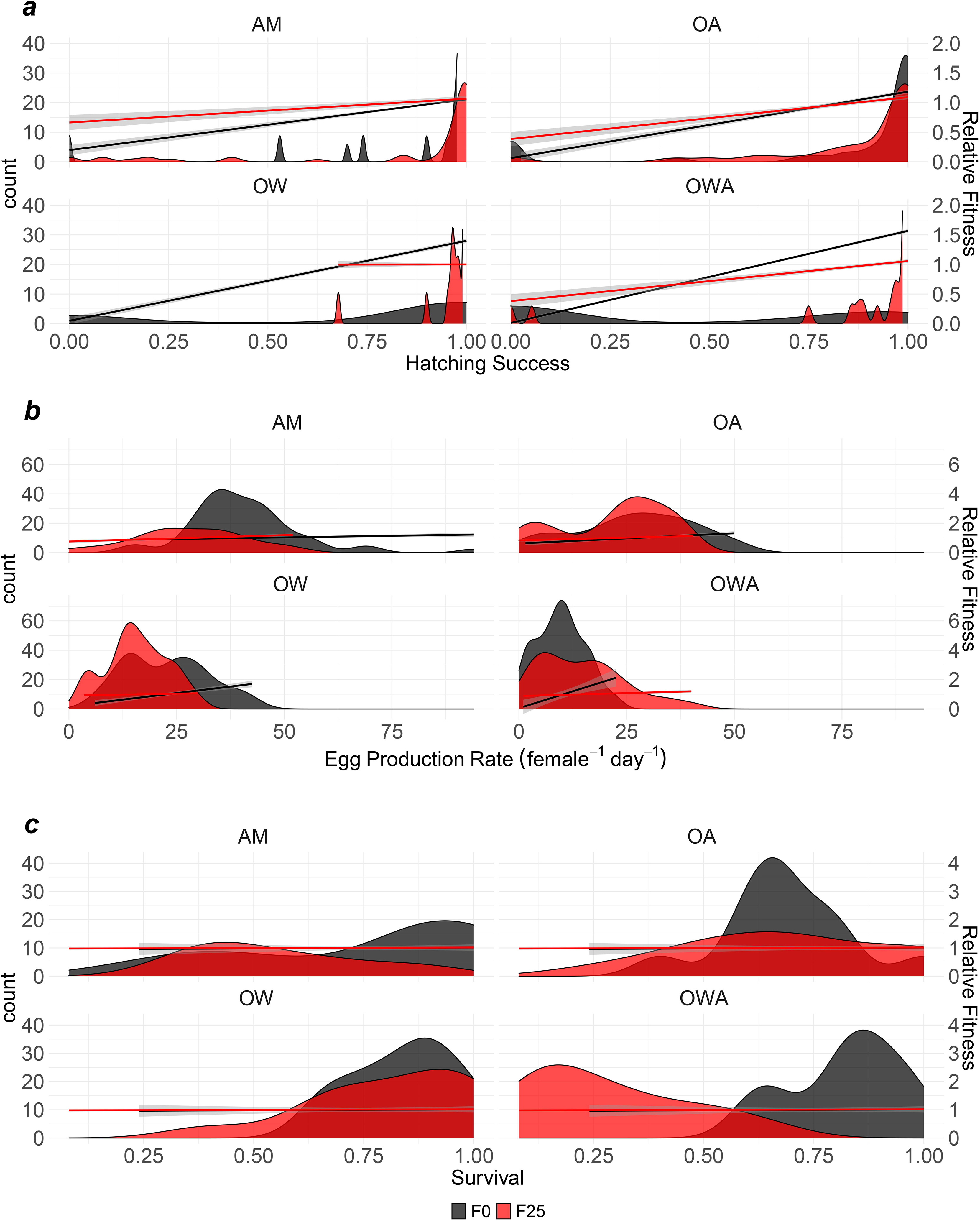
Fitness landscapes. Frequency distributions (left axes) and relative fitness landscape linear models (right axes) for **A**) Hatching success, **B**) Egg production rate, and **C**) Survival at F0 (black) and F25 (red). Fitness landscapes estimate the correlations between changing quantitative phenotypes and changing relative fitness. Shading around lines represents standard error.

We also employed path analysis on structural equation models (SEMs) to identify the hierarchical interactions of all life-history traits and the traits’ effects on fitness^43,44^. High correlation coefficients of SEMs indicate causality of a given trait’s effect towards fitness. The path analysis revealed that HS had the largest effect on fitness of all life-history traits across treatments, with the exception of the OW treatment at generation 25, and was significant at both generations 0 and 25 (Table 2). Taken together, we conclude that selection on HS both under OW and OWA conditions was the critical factor in adaptation to those environments.

**Table 2.** Path analysis and effects on lambda. Standardized parameter estimates from structural equation models for each trait’s effect on lambda at both F0 (gray bars) and F25 (white bars) for each treatment. The highest parameter estimate for each treatment at each generation is highlighted in bold.

|  | AM | OA | OW | OWA |
| --- | --- | --- | --- | --- |
| *Survival* | 0.12 | -0.09 | 0.03 | 0.08 |
|  | -0.09 | 0.13 | 0.72 | 0.33 |
| *EPR* | -0.07 | 0.04 | 0.03 | 0.10 |
|  | 0.34 | 0.25 | **0.86** | 0.23 |
| *HS* | **0.91** | **0.94** | **0.96** | **0.78** |
|  | **0.53** | **0.65** | 0.11 | **0.70** |
| *Sex ratio* | 0.12 | -0.15 | 0.09 | 0.21 |
|  | -0.39 | 0.37 | 0.37 | -0.27 |

## Implications

Our study highlights the need for global change studies to consider population fitness, in addition to individual traits, as an integrative tool for measuring adaptation^27,45^. For example, in this experiment using survival alone would have led to the erroneous conclusion of no adaptation under any treatment. Similarly, estimating fitness based on EPR alone would have also led to the erroneous conclusion that fitness decreased between generations 3 and 15 for AM. Despite this decrease in EPR, fitness remained unchanged across generations for AM (Fig. 3). Likewise, using EPR and HS alone would have missed the fitness decrease after generation 12 under OWA conditions. At the same time, the tradeoffs of EPR and HS versus survival after generation 12 (Figs. 1 and 2) partly explain the limited evolutionary rescue under OWA conditions (Fig. 3).

The shifting interactive effects of combined ocean warming and ocean acidification through time highlight the need to evaluate long-term evolutionary studies with multiple stressors and underline the complexity that accompanies predicting organism responses to climate change^33,46^. Previous work has shown that *Acartia tonsa* can adapt to lower CO_2_ levels (800 μatm) than those in our study^46^. Thus, it is conceivable that full evolutionary rescue to OWA can be achieved at lower CO_2_ levels. Furthermore, previous research has shown that exposure to OA over multiple generations improves EPR and selects for genes involved in RNA processing and regulation of metabolism in copepods^11,47^. Forthcoming genomic work from our group will determine if similar genes are also under selection for OW and OWA. Neither deleterious effects of OA nor adaptation to OA alone were observed across generations in our study (Fig. 3). Thus, our results highlight the evolutionary rescue and accompanying limitations that arise from multi-stressor adaptation. Assuming additive effects of warming and acidification in the present study should have led to full evolutionary rescue under OWA conditions, which was not observed (Fig. 3).

Instead, OW and OA had synergistic deleterious effects on fitness on generation 0, but antagonistic effects on generation 25 that counterbalance the individual effects of OW and OA (Supplementary Table 4). We suggest that the limited evolutionary rescue under OWA conditions in our study must have arisen from an antagonistic interaction between warming and acidification, as has been hypothesized earlier in polychaetes^19,20,48^ or for other co-occurring environmental stressors in bivalves^49^. Thus, since OW and OA are occurring simultaneously, even crude predictions of population performance under climate change should consider non-additive effects of temperature and CO_2_ interactions on population fitness.

Our study, showing both improved trait performance and population fitness across generations, constitutes the first demonstration of rapid adaptation to global change conditions for a metazoan. We present evidence for adaptation to both warming and combined warming and acidification, but not acidification alone, based on standing genetic variation. Previous work has been limited to phytoplankton^39^ or documenting marine metazoan traits for limited generations without population fitness estimates^18–20^. Failing to account for adaptation potential may overestimate future population vulnerability^45,50^. For example, using the observed deleterious effects at generation 0 to predict future vulnerability would have estimated a fitness reduction for the OWA treatment relative to ambient treatment of 56% (Fig. 3). However, accounting for evolutionary rescue (generations 3-25) resulted in an average fitness reduction of only 9%. At the same time, our results suggest that full evolutionary rescue under OWA conditions was not achieved (Fig. 3). This result is consistent with the hypothesis that adaptation to stress comes at a cost and that full recovery of populations to future climate conditions based on extant genetic variation is limited^51^, even though *A. tonsa* periodically experiences the warming and acidification conditions examined here^31,32^. The limited evolutionary rescue we observed also has consequences for future oceanic systems under OWA conditions. Because adaptation to OWA conditions appears to be costly even under the food replete conditions of our study, it is unclear to what extent evolutionary rescue would occur under scenarios that predict lower resource availability for zooplankton under climate change^50^. Limited evolutionary rescue, in turn, would reduce prey availability for fish, thereby negatively affecting fish production^50^. Finally, since copepods are a major vector of carbon transfer from the surface waters to the sea floor^28^, limited evolutionary rescue would also reduce the efficiency of the biological carbon pump, with a concomitant lower drawdown of CO_2_ from near surface waters.

## Conclusions

We show evidence for metazoan evolutionary adaptation to combined warming and acidification by explicitly assessing changes in population fitness based on a comprehensive suite of life-history traits. Adaptation was evident in the fitness improvements for animals after a few generations, and confirmed by independent evidence of genetic differentiation. The strength of selection coefficients and the path analysis suggested the key trait under selection affecting fitness in our study was hatching success. Evolutionary rescue was limited by the interaction of warming and acidification, which switched from synergistic to antagonistic from the beginning to the end of the experiment, adding complexity to predictions of population adaptation to climate change.

## Methods

### a) Copepod culturing and maintenance

Copepods were collected in June of 2016 from Esker Point Beach in Groton, CT, USA (41.320725°N, 72.001643°W) and raised for at least three generations as stock cultures prior to the start of transgenerational experiments to limit maternal effects^52^. Stock cultures were split evenly into eight groups of 160 females and 80 males. Four of these eight groups were acclimatized to high temperature at 1 degree Celsius per day and used to seed the two high temperature treatments (OW and OWA). The other four groups remained at ambient temperature and were used to seed the ambient and acidification treatments. After temperature acclimatization, groups of stock cultures seeded the parental (F0) individuals for two days. Stock culture groups yielded an average of 7,173 eggs per group to produce approximately 57,000 parental (F0) eggs. Resulting parental eggs and N1 nauplii were acclimated to one of four experimental treatments over the entire F0 generation: **1)** Ambient control (AM; temperature = 18°C, CO_2_ ~400 μatm; pH ~8.2); **2)** Ocean Acidification (OA; ambient temperature, high CO_2_ ~2000 μatm CO_2_, pH ~7.5); **3)** Ocean Warming (OW; high temperature = 22°C, ambient CO_2_); **4)** combined warming and acidification (OWA; high temperature, high CO_2_). Treatment replicates were derived from the stock culture groups (i.e. stock culture group 1 after high temperature acclimatization seeded the F0 for OW replicate 1 and OWA replicate 1). The ambient temperature was chosen from a Gaussian fit-model^53^ for optimal recruitment (egg production × egg hatching success) vs temperature derived from *Acartia tonsa* populations from Casco Bay (Gulf of Maine), Long Island Sound, and Chesapeake Bay (USA), 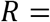 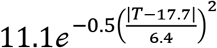 where R is recruitment (egg production rate × hatching success), T is temperature (°C), 17.7 is the optimal temperature and 6.4 is the standard deviation around the optimal temperature (N = 54, r^2^ = 0.42; our own unpublished data). Each treatment was kept in a separate temperature-controlled incubator (Thermo FisherScientific^®^ Isotemp™; Waltham, MA, USA) and split into four replicate 10L culture containers (Cambro; Huntington Beach, CA, USA). Copepods were fed every 48-72 h at food-replete concentrations (≥800 μg Carbon L^−1^) consisting of equal proportions of the phytoplankters *Tetraselmis* sp., *Rhodomonas* sp., and *Thalassiosira weissflogii*, following long-standing copepod culture protocols in our lab^54^. The phytoplankton fed to copepods was deliberately raised under ambient conditions for the entire length of the experiment to avoid confounding effects of possible changes in food quality due to the different temperature and CO_2_ among treatments. We minimized the chance of selecting for early developers in the cultures. Based on development time and adult longevity, we allowed copepods to contribute progeny to the next generation for 7-10 days once we observed the first nauplii of a new generation.

Elevated CO_2_ levels were achieved with gas proportioners (Cole-Parmer; Vernon Hills, IL, USA) mixing laboratory air with 100% bone dry CO_2_ that was delivered continuously to the bottom of each replicate culture. For small volume life-history trait experiments (sections b-d), CO_2_-mixed air was fed into custom plexiglass enclosures within each temperature-controlled incubator to allow for passive diffusion of CO_2_ into seawater. Target pH values were monitored using a handheld pH probe (Orion Ross Ultra pH/ATC Triode with Orion Star A121 pH Portable Meter (Thermo FisherScientific®; Waltham, MA, USA). The pH probe was calibrated monthly using commercially available NBS pH standards in a three-point calibration (pH 4.01, 7, 10.01; Thermo FisherScientific®; Waltham, MA, USA). Temperature and pH were monitored to ensure that small-volume experiments in the plexiglass enclosures matched those of bulk cultures. To counteract metabolic CO_2_ accumulation, control CO_2_ conditions were achieved by forcing compressed ambient air through a series of CO_2_-stripping units containing granular soda lime (AirGas®; Waterford, CT, USA) and a particle filter (1 μm), and then to each culture container via airstone. Continuous bubbling maintained dissolved oxygen levels at >8 mg L^−1^. Temperature, pH, and actual *p*CO_2_ were monitored throughout the experiment (Supplementary Table 1). Actual *p*CO_2_ conditions were calculated in CO_2_SYS^55^ based on measurements of salinity, temperature, pH, and total alkalinity (*A*_T_; Supplementary Tables 2-3) with k1/k2 from Lueker, et al. 2000 ^56^, KHSO_4_ from Dickson 1990 ^57^, total Boron from Uppstrom 1974 ^58^, and pH based on NBS scale. Total alkalinity was measured in triplicates three times over the course of the experiment using endpoint titration (G20 Potentiometric Titrator; Mettler Toledo^®^)^59^.

Because treatments were housed in separate incubators, incubator-specific effects are theoretically possible, but unlikely^60,61^ given that incubators were held under identical ambient conditions except for the constant temperature and CO_2_ conditions that were meticulously monitored, and verified by independent measures throughout the experiment (Supplementary Table 5). Because of logistical and personnel constraints, egg production rate (EPR), hatching success (HS), survival, and development time (see below) could not be measured at every generation. Instead, measurements were taken at generations 0, 3, 6, 9, 12, 15, and 25.

### b) Egg production rate and hatching success

For each replicate culture within a treatment, 10 pairs of newly developed males and females were placed into 20 mL petri dishes for 48 h (n = 280 per treatment). The dishes were housed in custom-made, airtight, plexiglass enclosures whose atmosphere was controlled to the appropriate CO_2_ concentration (see *copepod culturing and maintenance* section). There was one enclosure per temperature-controlled incubator. After the 48-h egg laying period, adults were checked for survival and removed from the petri dishes. Eggs were left in the dishes for an additional 72 h to allow for egg hatching and their contents preserved with non-acid Lugol’s solution. Dishes with dead males were used for EPR, but not HS, since fertilization could not be assumed. Dishes with dead females were discarded. EPR was calculated as 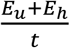 where *E_u_* represents unhatched eggs, *E_h_* represents hatched eggs (nauplii), and *t* represents egg laying time. Hatching success was calculated as 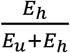.

### c) Survival

Survival was measured from nauplius 1 (N1) stage to copepodid 6 (adult) stage. For a given generation, all adults from the previous generation were removed from the culture and allowed to lay eggs in food-replete media for 48 h. Resulting nauplii were chosen for tracking survival. Unhatched eggs and any nauplii not chosen for survival analysis were returned to their respective replicate cultures for continued population maintenance. To measure survival for all generations where life-history traits were evaluated, three 250-mL beakers for each replicate culture were supplied with 25 randomly chosen N1 nauplii each and housed in the plexiglass enclosure described above (n = 21 per treatment). Copepods were checked every 48-72 h. The number of dead, live, and missing copepods were logged for each beaker along with general stage (i.e. nauplius, copepodite, adult female, or adult male). The fraction of survived individuals (*l*_*x*_) was calculated as *n*_*x*_/*n_i_* where *n_x_* represents the number of live individuals on day *x*, and *n_i_* represents initial individuals. Nauplii were grown with media at levels of 500 μg C L^−1^ to prevent overgrowth of phytoplankton and allow for adequate nauplii grazing. Following the naupliar stages, copepods were grown with food-replete (800 μg C L^−1^) media as described earlier. Food was replaced with fresh media on monitoring days. Average survival was calculated per each replicate culture at each generation measured. Differences in day-specific survival between replicates and treatments was assessed using the ‘survival’ package in R^62^.

### d) Development time

To calculate development time (time from N1 to adulthood) we recorded the number of days at which adults matured during the survival experiments and averaged the observations.

### e) Sex ratio

Sex ratio was calculated based on the number of surviving adult females relative to surviving adult males in survival experiments.

### f) Fitness

The population net reproductive rate, λ, was calculated as the dominant eigenvalue of an assembled projected age-structured Leslie Matrix constructed from survival and fecundity data^37^. Briefly, day-specific probabilities of survival are calculated from day-specific survival as 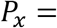 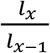 where *l_x_* represents the proportion of individuals on day *x* and *l_x−1_* represents the proportion of individuals on day *x−1*. Probabilities of survival on day 1 are assumed to be 100% or a value of “1.0”. Per capita EPR and HS are calculated as described above, with fecundity rates equaling the product of EPR and HS. Because only females produce offspring, total fecundity rates must be scaled to the sex ratio (proportion of females:males). To account for differences in individual development time for each treatment, fecundity rates are assigned to all days after the first matured adult is observed. We assume that surviving individuals represented by the survival experiments are equally as likely to experience any of the fecundity values observed in EPR experiments. Therefore, each mate-pair fecundity rate was paired with each survival beaker to construct a matrix. This yields a maximum of 120 matrices per treatment per generation (3 survival beakers × 4 replicate cultures × 10 mate pairs).

### g) Selection coefficients and path analysis

Linear selection coefficients (β) were evaluated by creating multiple linear regression models of relative fitness 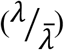 against survival, egg production rate, hatching success, and sex ratio at both F0 and F25 for each treatment and calculating the partial regression coefficient estimate for each trait^63^. The path analysis was completed by creating structural equation models^43,44^ of relative fitness against survival, egg production rate, hatching success, and sex ratio at both F0 and F25 for each treatment using the ‘lavaan’ and ‘semPlot’ packages in R^64,65^. Development time was omitted from the models because the lack of variance within a replicate violated assumptions of the model.

### h) Genetic Diversity

To quantify genetic diversity across the genome, capture probes were designed to target both coding and regulatory regions across the genome. For coding regions, we chose the highest quality probe falling within the region. Regulatory probes were set within 1,000 bp upstream of the transcription start site. Genomic DNA was shipped to Rapid Genomics (Gainesvill, FL, USA) for library preparation and was captured with the 32,413 probes (21,311 coding, 11,102 regulatory). Enriched libraries were sequenced on a HiSeq 4000 with 150 bp paired end reads. Raw data were trimmed for quality and adapter contamination with Trimmomatic v0.36^66^. Trimmed reads were mapped to the *A. tonsa* reference genome^67^ with BWA-MEM^68^.

SAMTOOLs^69^ was used to generate a pileup for each sample, from which genetic diversity (*π*) was estimated with Popoolation^70^. We identified 1,450 100bp windows in 704 unique scaffolds across the genome that were present across all samples. We estimated *π* in 100bp sliding windows with a 100bp step size. Each position required a minimum coverage of 30×, max coverage of 1000× (to avoid mapping errors), and at least 0.5 of the window meeting these thresholds. Resulting windows were required to be sequenced across all samples. To take into account the independent replicates within each treatment, we used pairwise Wilcoxon Rank Sum tests with a Holm correction for multiple testing. Genomic data is deposited in GenBank: BioProject number PRJNA590963. All statistics were performed in R^71^.

### i) Statistical analyses

Statistical analyses were computed using the software package R (v 4.0.2)^71^. To examine effects of generation on changing life-history traits, we created trait-specific generalized additive models (GAMs) smoothed across generations for each treatment^72,73^. To evaluate differences between life-history traits, we created separate linear mixed models with replicates as random effects and used post-hoc t-tests and Tukey HSD tests to compare life-history trait values that were significantly different from other treatments at each generation (α < 0.05). Analysis of fitness (λ) calculations also included estimations with a zero-inflated generalized linear mixed effects model with generation and treatment as fixed effects and replicates as random effects. Analyzing the data with linear mixed models also allowed us to evaluate the effects of treatment replicates on life-history traits. A low intra-class correlation coefficient suggests no predictive effect of random variables. For the model constructed for fitness over generations, the variance due to replicates within a treatment is very low (ICC = 0.08; Table 1) and does not affect the model results. To estimate the predicted probabilities of λ = 0 across generations in Supplementary Fig. 4, we converted λ values to either 0 or 1 representing λ values of 0 and >0, respectively. We used the binomial-converted λ data to fit a linear mixed effects model against generation and treatment with replicates included as random effects. To evaluate individual effects of temperature, pH, or generation on life-history traits, we constructed a third linear model that was tested with a three-way analysis of variance.

## Acknowledgements

Research supported by grants from the USA National Science Foundation (OCE-1559180 awarded to HGD, MBF and HB and OCE-1559075 awarded to MHP) and Connecticut Sea Grant (R/LR-25) awarded to HGD, MBF, and HB.. The authors thank W. Huffman for aiding in pilot experiments, C. Murray for assistance in alkalinity measurements, D. Arbige, C. Woods, and B. Dziomba for help in maintaining equipment and constructing custom enclosures for the experiments, and T. Moore and J. Lee of UConn’s Statistical Consulting Services for advice and assistance on data analysis.

## Author Contributions

HGD conceived the project, designed research, aided in data analysis, and wrote the manuscript. JAD conducted experiments, analyzed data, created figures, and wrote the manuscript with HGD. GP, LN, and XH conducted experiments. MBF conceived the project and designed research. HB conceived the project, designed research, and designed the CO_2_ delivery system. RSB performed genomic diversity analysis. MHP conceived the project, designed research, and performed genomic analysis. All authors edited and approved the paper.

## Data and Script availability

The phenotypic data and R-scripts referred in the text have been deposited in zenodo (https://doi.org/10.5281/zenodo.4477401). The genetic diversity data are deposited in GenBank: BioProject number PRJNA590963, to be released upon publication of this paper.

