## Supplementary for "Rapid, but limited, zooplankton adaptation to simultaneous warming and acidification"

**Supplementary information for Dam et al. Rapid, but limited, zooplankton adaptation to simultaneous warming and acidification**

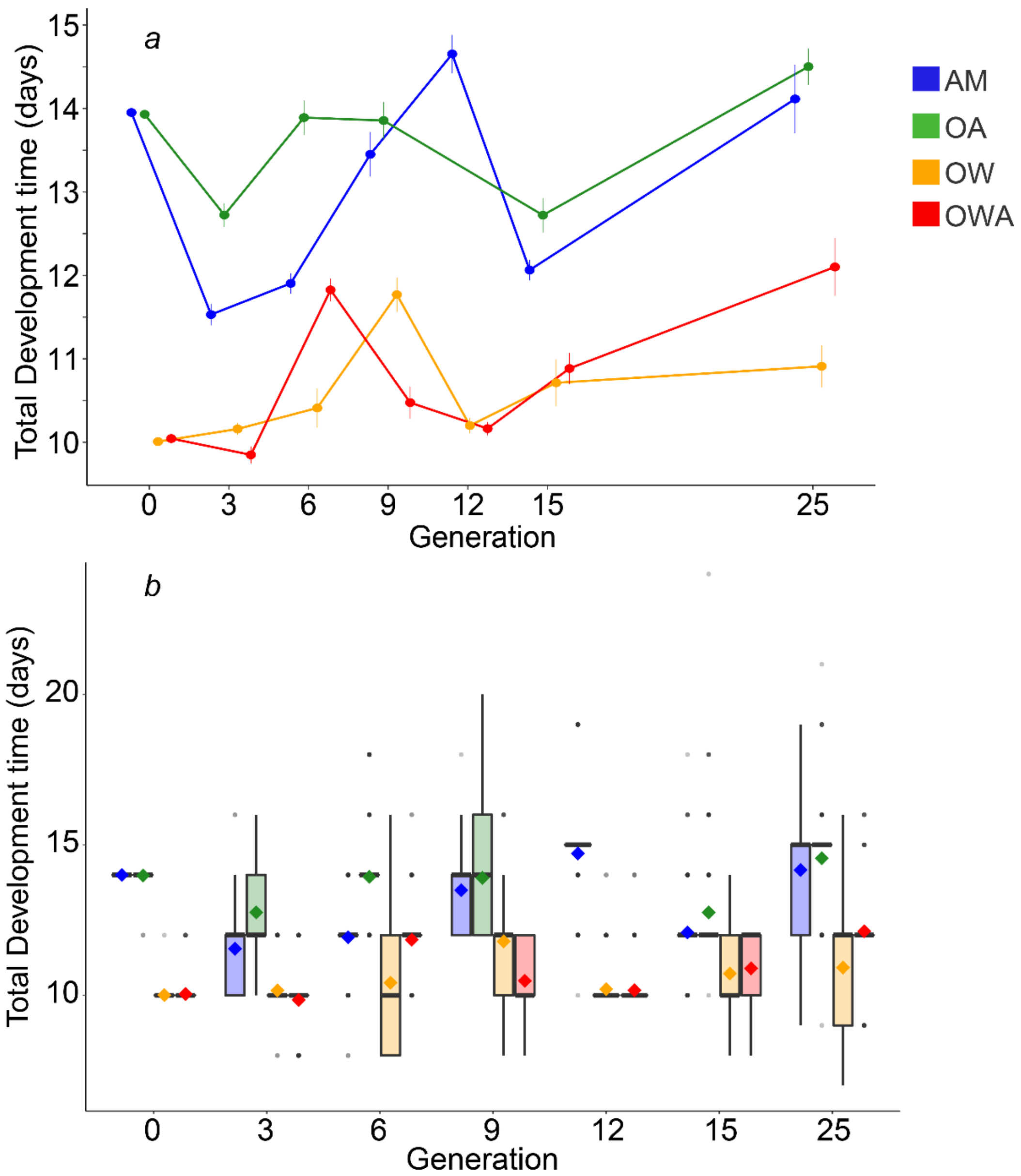


***Supplementary Figure 1 – Development time for transgenerational study.*** *A) Mean development time across generations. Error bars represent 95% confidence intervals. B) Box and whisker plots displaying data distribution. In the boxes, the center black line represents the median, diamonds represent the mean, upper box edge represents the 75% quartile, lower box edge represents 25% quartile, whiskers represent 1.5x interquartile range, and points represent outliers. Curves for treatments are offset for clarity in panel A. Treatment colors: blue: AM; green: OA; orange: OW; red: OWA.*

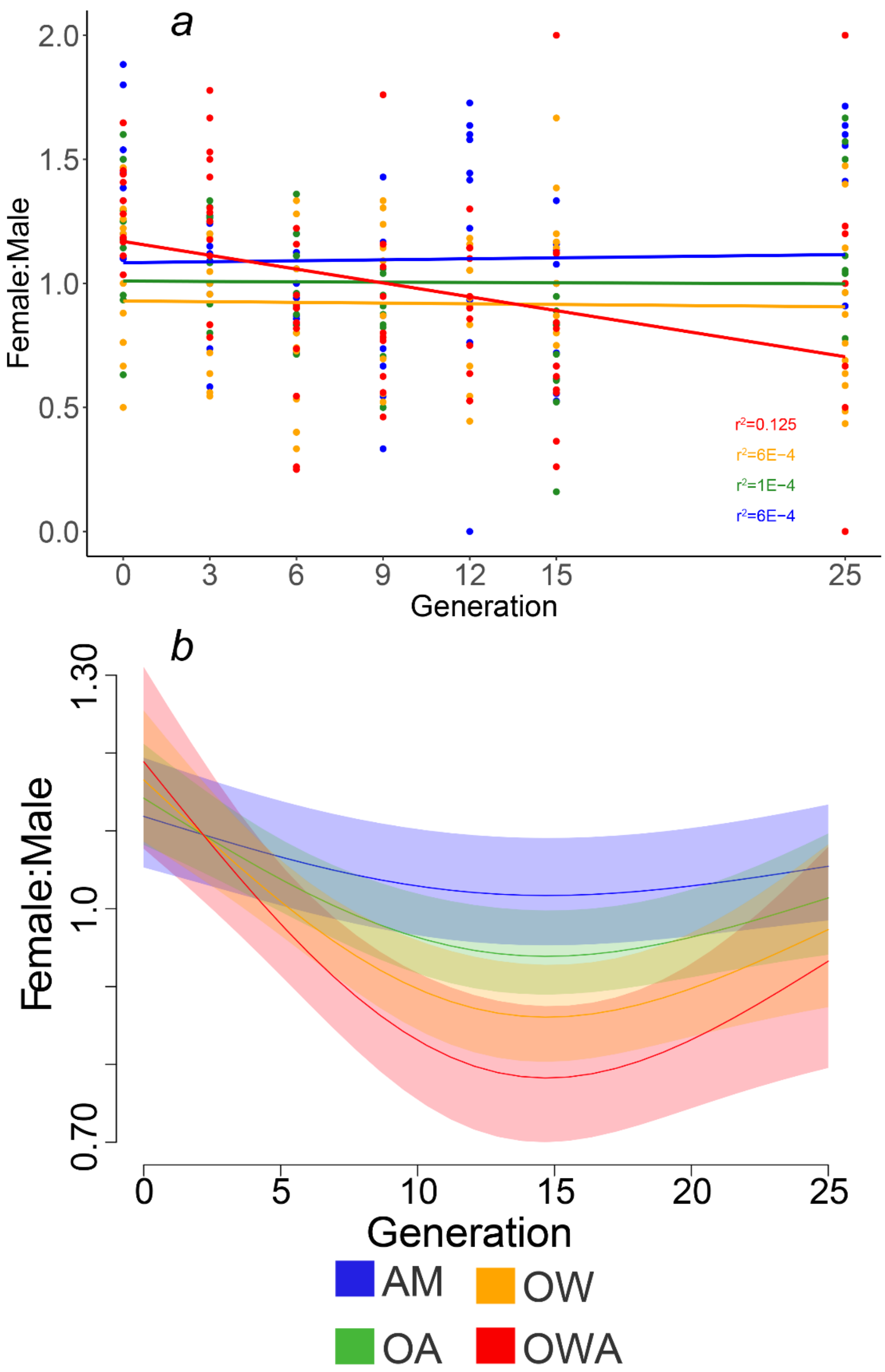


***Supplementary Figure 2 – Sex ratio for transgenerational study.*** *Results for sex ratio across generations modeled as A) linear model and B) GAM. Treatment colors: blue: AM; green: OA; orange: OW; red: OWA.*

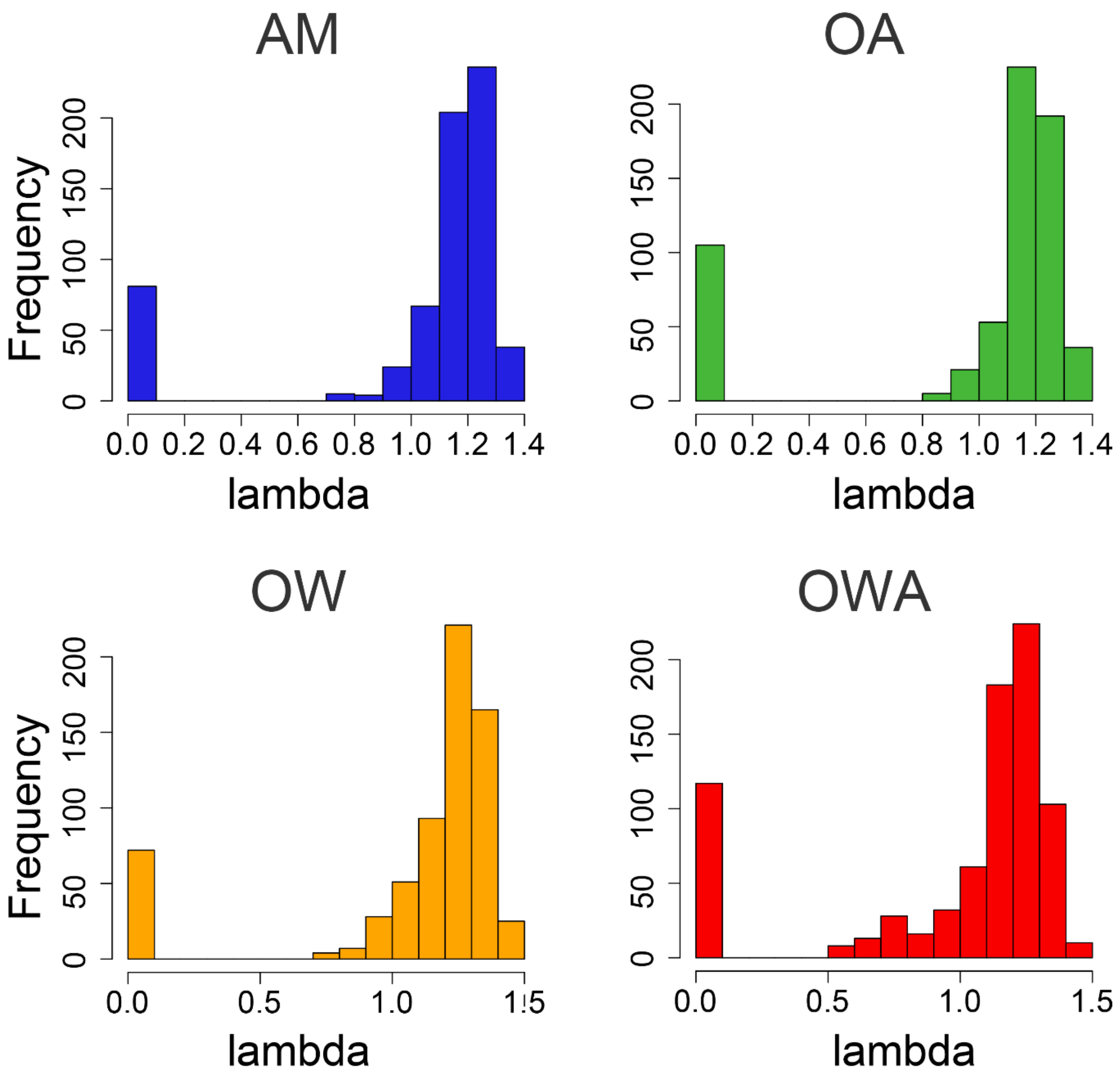


***Supplementary Figure 3 – Histogram of fitness values.*** *Frequency of lambda (λ) values for all generations by treatment. Treatment colors: blue: AM; green: OA; orange: OW; red: OWA.*

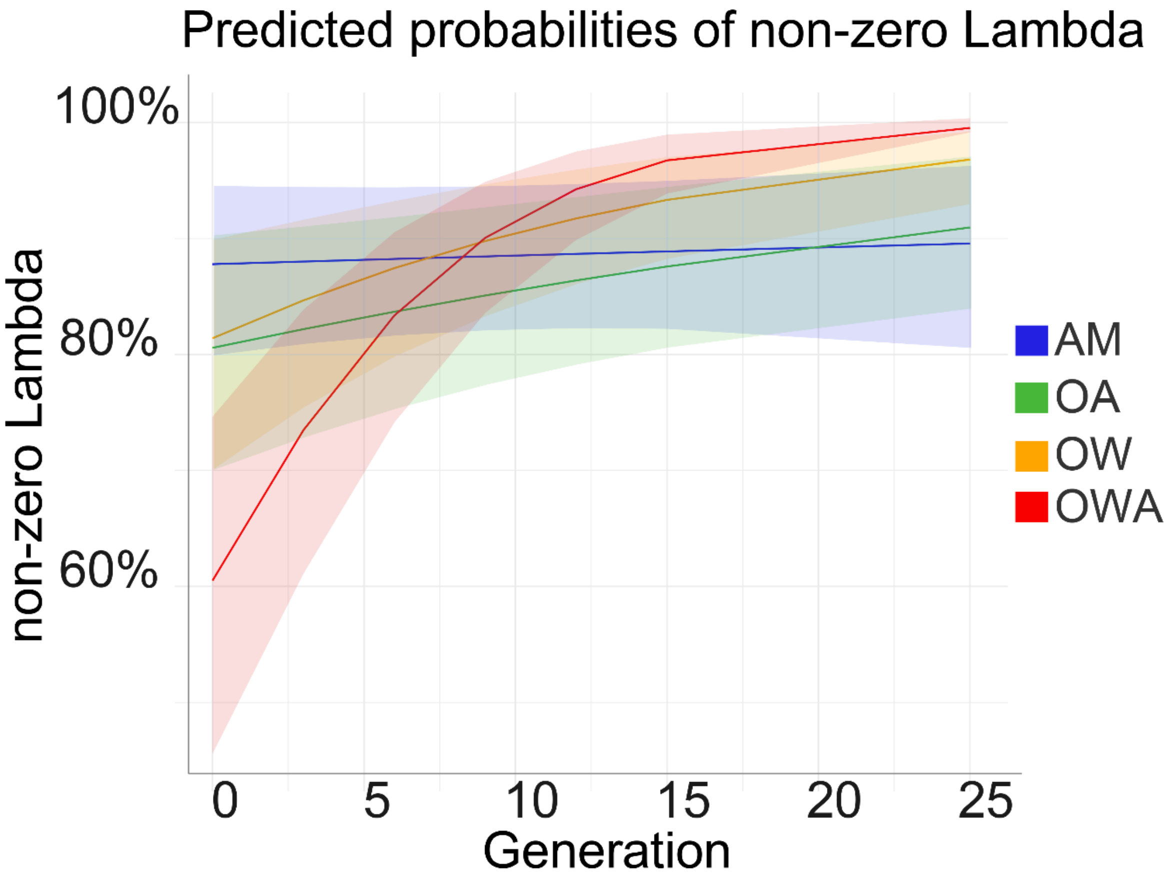


***Supplementary Figure 4 – Predicted probabilities of non-zero lambda.*** *Predicted probabilities of lambda that is not zero. Probabilities for ambient, acidification, and warming treatments are unchanging across generations. Probability of Greenhouse non-zero lambda values increase over time. Shading represents 95% confidence intervals. Treatment colors: blue: AM; green: OA; orange: OW; red: OWA.*

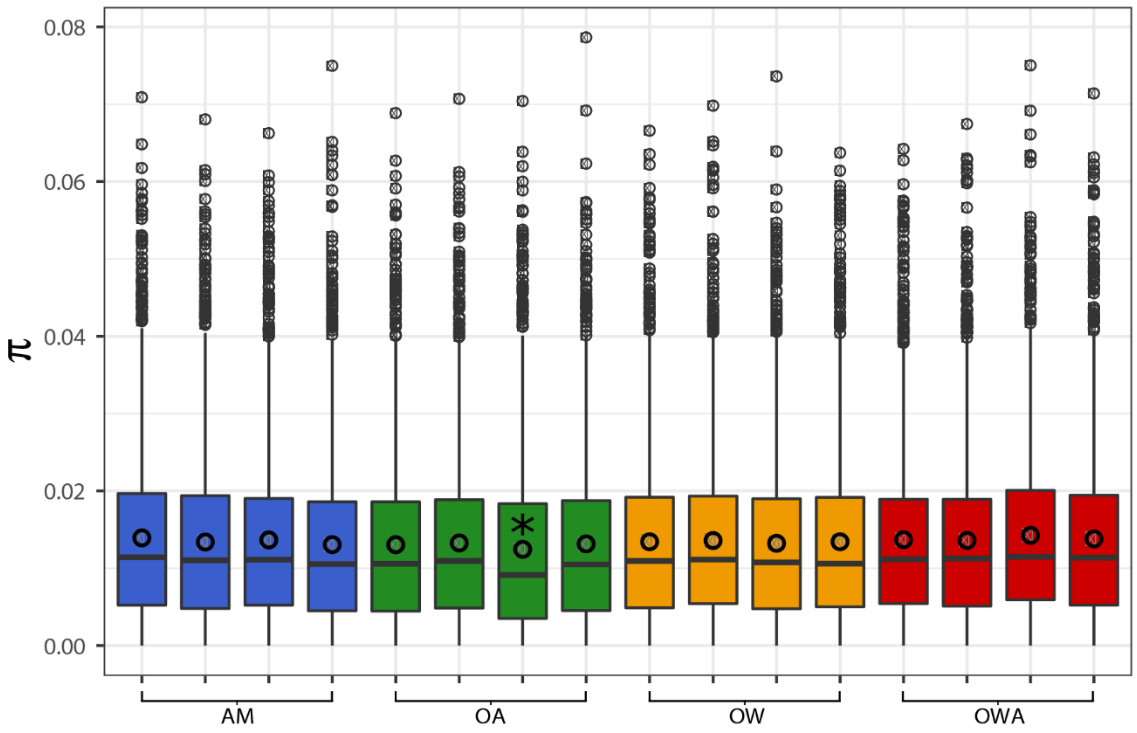


***Supplementary Figure 5 – Genetic Diversity.*** *Estimates of genetic diversity (π) after 25 generations of lab conditions. Estimates were calculated in 100 bp non-overlapping sliding windows. Windows were included when at least 50% of sites had coverage between 30x and 1000x per sample and the window was covered across all samples. The asterisk indicates the Acidification sample with reduced genetic diversity relative to other samples (Wilcoxon Rank Sum test with Holm correction for multiple testing; $p < 0.05$); all other samples were not significantly different ($p > 0.05$). In the boxes, the center black line represents the median, the circles represent means, upper box edge represents the 75% quartile, lower box edge represents 25% quartile, whiskers represent 1.5x interquartile range, and points represent outliers. Treatment colors: blue: AM; green: OA; orange: OW; red: OWA.*

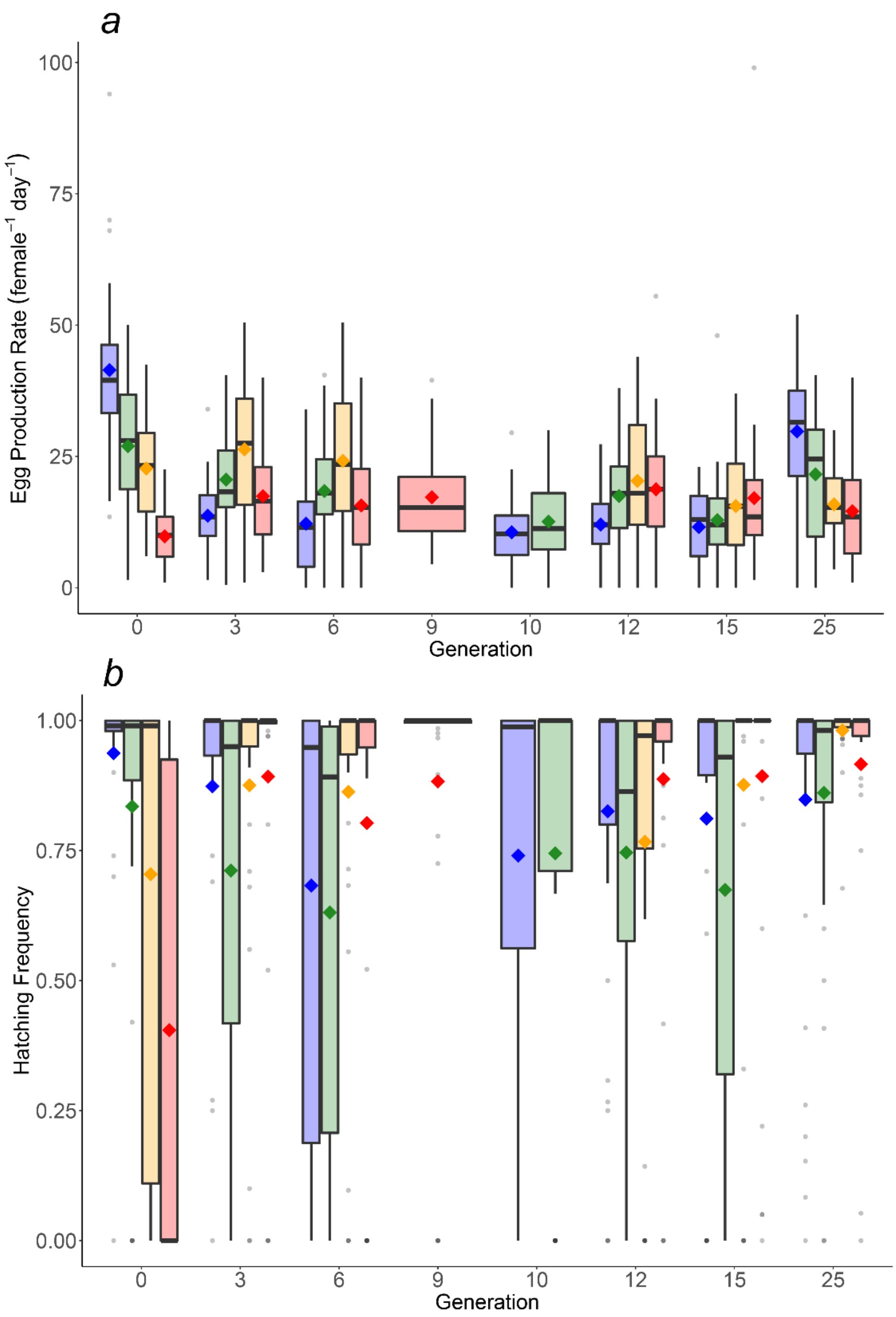


***Supplementary Figure 6 – Box and whisker plots for figure 1 in the text.*** *A) Egg production rate and B) Hatching frequency. In the boxes, the center black line represents the median, diamonds represent the mean, box edges represent the 25 and 75% quartiles, whiskers represent 1.5x interquartile range, and points represent outliers. Treatment colors: blue: AM; green: OA; orange: OW; red: OWA.*

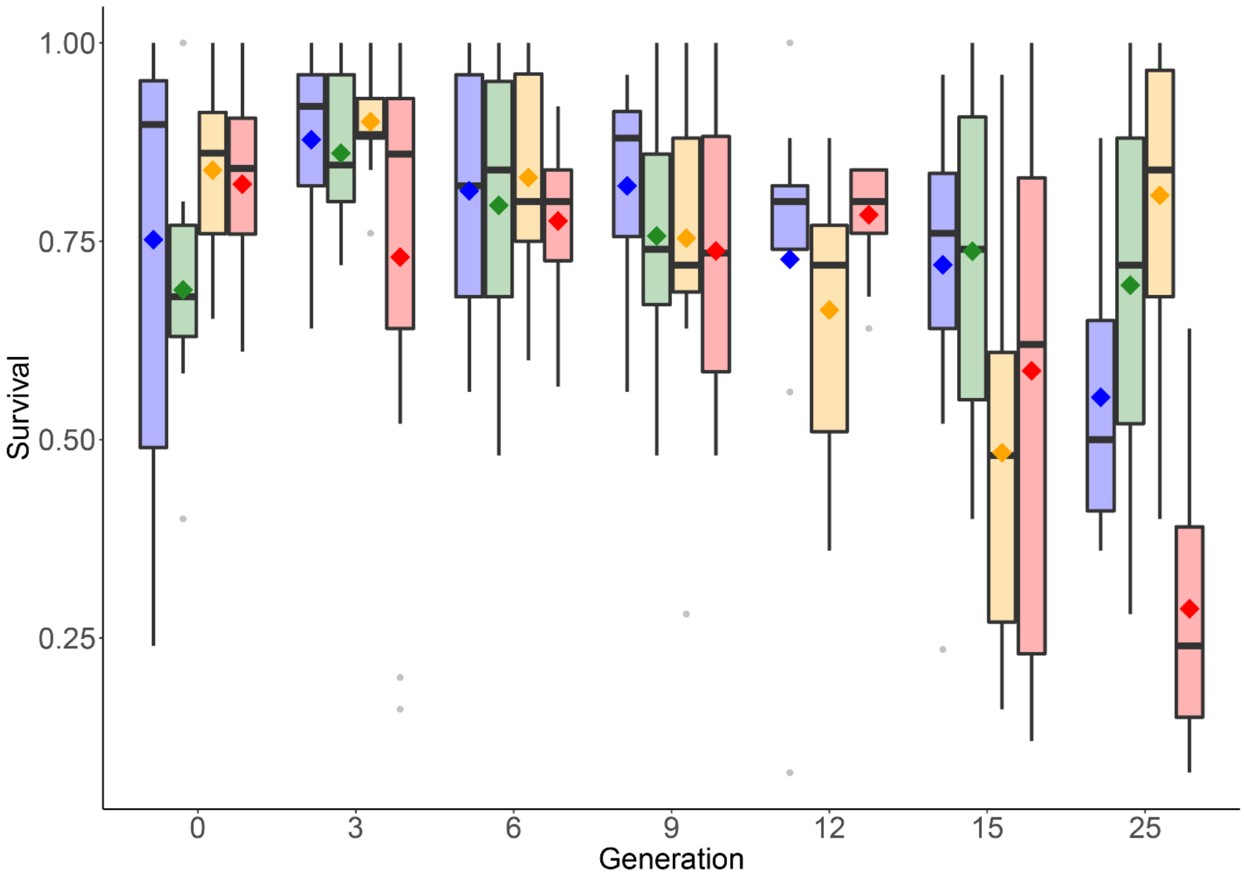


***Supplementary Figure 7 – Box and whisker plots for figure 2 in the text (survival)****. In the boxes, the center black line represents the median, the diamonds represent the mean, box edges represent the 25 and 75% quartiles, whiskers represent 1.5x interquartile range, and points represent outliers. Treatment colors: blue: AM; green: OA; orange: OW; red: OWA.*

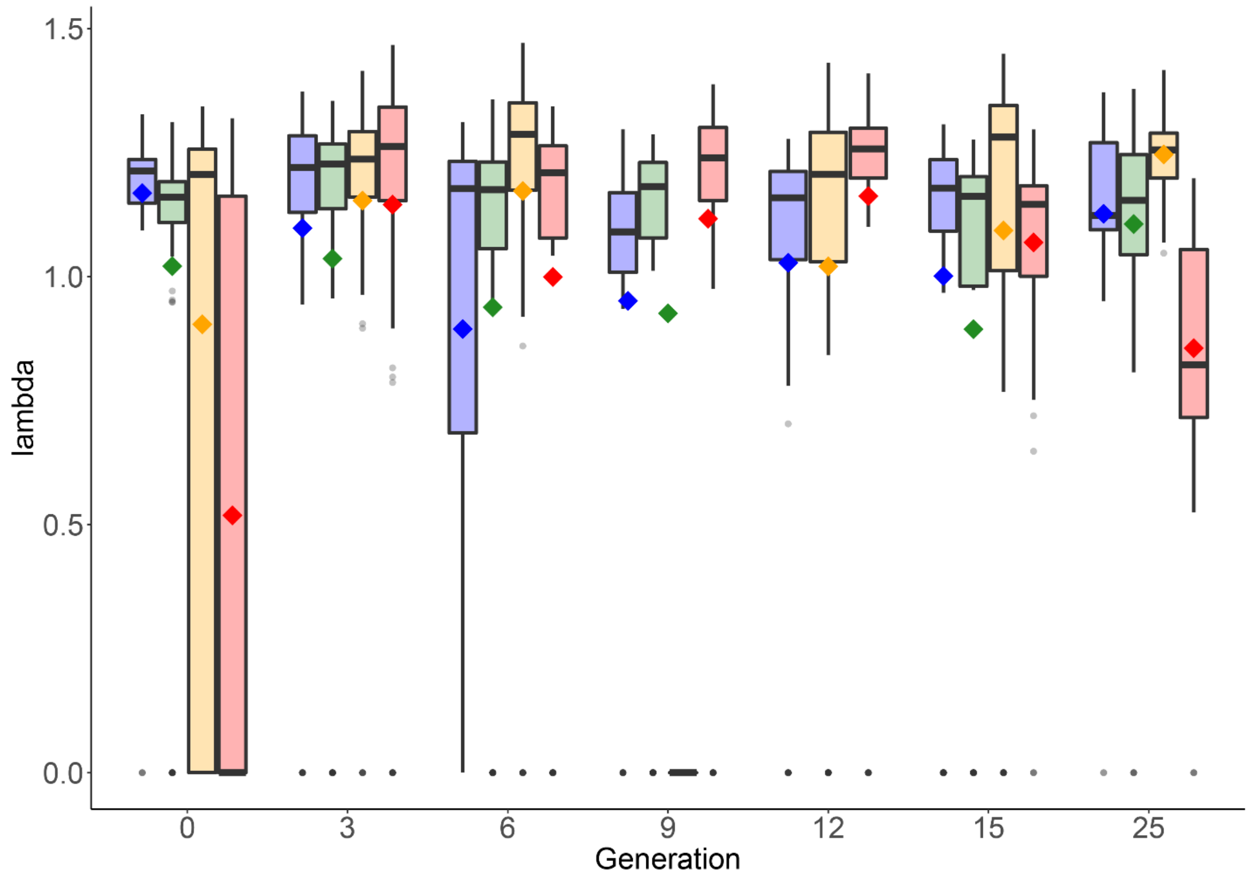


***Supplementary Figure 8 – Box and whisker plots for Figure 3 in the text (Fitness).*** *In the boxes, the center black line represents the median, the diamonds represent the mean, box edges represent the 25 and 75% quartiles, whiskers represent 1.5x interquartile range, and points represent outliers. Treatment colors: blue: AM; green: OA; orange: OW; red: OWA.*

***Supplementary Table 1 – Temperature and $CO_2$ measurements.*** *Mean temperature, pH, and $pCO_2$ measurements collected across the transgenerational experiment. SD = standard deviation. N = number of observations. SE = standard error.*

| Variable | Treatment | Target value | Mean measured value | SD | N | SE |
|---|---|---|---|---|---|---|
| **Temperature** | AM | 18°C | 18.08 | 0.36831 | 165 | 0.028673 |
| | OA | 18°C | 17.95 | 0.302756 | 165 | 0.02357 |
| | OW | 22°C | 21.90 | 1.098083 | 167 | 0.084972 |
| | OWA | 22°C | 21.94 | 0.334943 | 169 | 0.025765 |
| **pH** | AM | 8.2 | 8.28 | 0.10119 | 165 | 0.007878 |
| | OA | 7.5 | 7.53 | 0.082598 | 165 | 0.00643 |
| | OW | 8.2 | 8.25 | 0.104003 | 167 | 0.008048 |
| | OWA | 7.5 | 7.57 | 0.079519 | 169 | 0.006117 |
| **$pCO_2$** | AM | 400 μatm | 380.94 | 49.29688 | 9 | 16.43229 |
| | OA | 2000 μatm | 2245.09 | 276.6075 | 9 | 95.20249 |
| | OW | 400 μatm | 377.17 | 17.67343 | 9 | 5.891142 |
| | OWA | 2000 μatm | 2358.12 | 120.8067 | 9 | 40.26889 |

***Supplementary Table 2 – Total alkalinity and pCO$_2$ measurements.*** *Mean ± (SD; SE) of total alkalinity ($A_T$; µmol kg$^{-1}$), partial pressure of $CO_2$ (pCO$_2$; µatm), fugacity of $CO_2$ (fCO$_2$; µatm), $\Omega_{Ca}$, $\Omega_{Ar}$, total DIC (µmol kg$^{-1}$), and salinity measured from replicate measurements of seawater samples taken during the transgenerational experiment. Salinity was measured via refractometer and $A_T$ from endpoint titrations. N represents the number of observations. Variables calculated by $CO_2$SYS (pCO$_2$ and fCO$_2$) are denoted by an asterisk (*). Variables calculated with k1/k2 constants from Leuker, et al. 2000 [54] and $KHSO_4$ from Dickson, 1990 [55], total Boron from Uppstrom 1974 [56], and pH based on NBS scale.*

| Measurement | Temperature (°C) | pH | Target *p*CO$_2$ | Salinity | $A_T$ | *p*CO$_2$* | fCO$_2$* | N |
|---|---|---|---|---|---|---|---|---|
| 1 | 18 | 8.2 | 400 | 31 | 2086 ± (10; 6) | 348 ± (2; 1) | 347 ± (2; 1) | 3 |
| | | 7.5 | 2000 | 31 | 2106 ± (10; 6) | 2062 ± (10; 6) | 2056 ± (10; 6) | 3 |
| | 22 | 8.2 | 400 | 31 | 2146 ± (23; 13) | 366 ± (4; 2) | 364 ± (4; 2) | 3 |
| | | 7.5 | 2000 | 31 | 2170 ± (10; 6) | 2209 ± (10; 6) | 2201 ± (10; 6) | 3 |
| 2 | 18 | 8.2 | 400 | 36 | 2830 ± (119; 69) | 446 ± (10; 6) | 445 ± (10; 6) | 3 |
| | | 7.5 | 2000 | 35 | 2748 ± (19; 15) | 2613 ± (31; 17) | 2604 ± (30; 17) | 3 |
| | 22 | 8.5 | 400 | 33 | 2401 ± (27; 15) | 400 ± (5; 3) | 399 ± (5; 3) | 3 |
| | | 7.5 | 2000 | 35 | 2466 ± (56; 33) | 2441 ± (66; 38) | 2433 ± (66; 38) | 3 |
| 3 | 18 | 8.2 | 400 | 31 | 2095 ± (6; 3) | 348 ± (3; 2) | 348 ± (3; 2) | 3 |
| | | 7.5 | 2000 | 31 | 2103 ± (6; 3) | 2059 ± (6; 3) | 2052 ± (6; 3) | 3 |
| | 22 | 8.2 | 400 | 33 | 2200 ± (11; 6) | 366 ± (2; 1) | 364 ± (2; 1) | 3 |
| | | 7.5 | 2000 | 34 | 2443 ± (62; 36) | 2422 ± (60; 35) | 2416 ± (60; 34) | 3 |

***Supplementary Table 3 – Carbonate measurements.*** *Mean ± (SD; SE) of $\Omega_{Ca}$, $\Omega_{Ar}$, and total DIC (μmol kg$^{-1}$), calculated from temperature, pH, salinity and $A_T$ measured from replicate measurements of seawater samples taken during the transgenerational experiment. Salinity was measured via refractometer and $A_T$ from endpoint titrations. N represents the number of observations. Variables calculated by $CO_2$SYS ($\Omega_{Ca}$, $\Omega_{Ar}$, and total DIC) are denoted by an asterisk (*). Variables calculated with k1/k2 constants from Leuker, et al. 2000 [54], $KHSO_4$ from Dickson, 1990 [55], total Boron from Uppstrom 1974 [56], and pH based on NBS scale.*

| Measurement | Temperature (°C) | pH | Target $pCO_2$ | Salinity | $\Omega_{Ar}$* | $\Omega_{Ca}$* | DIC* | N |
|---|---|---|---|---|---|---|---|---|
| **1** | 18 | 8.2 | 400 | 31 | 2.4 ± (0.2; 0.009) | 3.8 ± (0.02; 0.01) | 1872 ± (9; 5) | 3 |
| | | 7.5 | 2000 | 31 | 0.57 ± (0.006; 0.003) | 0.88 ± (0.006; 0.003) | 2125 ± (10; 6) | 3 |
| | 22 | 8.2 | 400 | 31 | 2.8 ± (0.04; 0.02) | 4.3 ± (0.05; 0.03) | 1902 ± (21; 12) | 3 |
| | | 7.5 | 2000 | 31 | 0.68 ± (0.006; 0.003) | 1.0 ± (0.006; 0.003) | 2178 ± (10; 6) | 3 |
| **2** | 18 | 8.2 | 400 | 36 | 3.6 ± (0.3; 0.1) | 5.6 ± (0.4; 0.2) | 2516 ± (92; 53) | 3 |
| | | 7.5 | 2000 | 35 | 0.81 ± (0.01; 0.007) | 1.2 ± (0.02; 0.01) | 2764 ± (20; 12) | 3 |
| | 22 | 8.5 | 400 | 33 | 3.3 ± (0.04; 0.02) | 5.0 ± (0.06; 0.03) | 2121 ± (25; 14) | 3 |
| | | 7.5 | 2000 | 35 | 0.82 ± (0.02; 0.009) | 1.3 ± (0.02; 0.009) | 2466 ± (59; 33) | 3 |
| **3** | 18 | 8.2 | 400 | 31 | 2.4 ± (0.02; 0.01) | 3.8 ± (0.03; 0.02) | 1877 ± (9; 5) | 3 |
| | | 7.5 | 2000 | 31 | 0.57 ± (0.006; 0.003) | 0.89 ± (0.006; 0.003) | 2121 ± (6; 3) | 3 |
| | 22 | 8.2 | 400 | 33 | 3.0 ± (0.02; 0.009) | 4.6 ± (0.02; 0.01) | 1936 ± (10; 6) | 3 |
| | | 7.5 | 2000 | 34 | 0.81 ± (0.03; 0.02) | 1.2 ± (0.04; 0.02) | 2444 ± (62; 36) | 3 |

| Generation | Treatment | Mean λ value | SD | N | SE | λ Direction (Interaction) |
|---|---|---|---|---|---|---|
| **0** | AM | 1.168196 | 0.205603 | 107 | 0.019876 | |
| | OA | 1.021074 | 0.384388 | 100 | 0.038439 | Decrease |
| | OW | 0.903853 | 0.547074 | 102 | 0.054168 | Decrease |
| | OWA | 0.51869 | 0.585064 | 108 | 0.056298 | Decrease (*Synergistic*) |
| **3** | AM | 1.09774 | 0.378392 | 90 | 0.039886 | |
| | OA | 1.036184 | 0.454502 | 96 | 0.046387 | No change |
| | OW | 1.153172 | 0.304468 | 105 | 0.029713 | No change |
| | OWA | 1.145091 | 0.373395 | 108 | 0.03593 | No change (*Antagonistic)* |
| **6** | AM | 0.89433 | 0.523122 | 120 | 0.047754 | |
| | OA | 0.938287 | 0.491192 | 100 | 0.049119 | No change |
| | OW | 1.172572 | 0.357428 | 120 | 0.032629 | Increase |
| | OWA | 0.99928 | 0.467307 | 120 | 0.042659 | No change (*Antagonistic*) |
| **9** | AM | 0.951109 | 0.399669 | 70 | 0.04777 | |
| | OA | 0.925874 | 0.503452 | 120 | 0.045959 | No change |
| | OWA | 1.116796 | 0.381839 | 120 | 0.034857 | No information |
| **12** | AM | 1.027817 | 0.348379 | 100 | 0.034838 | |
| | OW | 1.020707 | 0.455653 | 117 | 0.042125 | No change |
| | OWA | 1.162178 | 0.338253 | 120 | 0.030878 | No information |
| **15** | AM | 1.001211 | 0.440747 | 114 | 0.04128 | |
| | OA | 0.893952 | 0.499696 | 111 | 0.047429 | Decrease |
| | OW | 1.093036 | 0.402368 | 120 | 0.036731 | No change |
| | OWA | 1.068935 | 0.217377 | 120 | 0.019844 | No change (*Antagonistic*) |
| **25** | AM | 1.126585 | 0.241798 | 58 | 0.03175 | |
| | OA | 1.105866 | 0.253182 | 110 | 0.02414 | No change |
| | OW | 1.246251 | 0.073536 | 102 | 0.007281 | Increase |
| | OWA | 0.855734 | 0.244107 | 99 | 0.024534 | Decrease (*Antagonistic*) |

***Supplementary Table 4 – Synergistic and antagonistic effects of temperature and $CO_2$ on fitness.*** *The direction of influence (statistically significant increase or decrease) relative to control is summarized for λ at each generation with the observed synergistic or antagonistic effect of the two environmental variables highlighted at the same time point in parentheses. No information on synergy or antagonism can be inferred for generations 9 and 12 because we are missing fitness data for one of the two treatments at each generation. SD = standard deviation, N = number of observations, SE = standard error.*

***Supplementary Table 5 – Contrasts of temperature, pH, and $pCO_2$ across treatments.*** *Comparison of physical environmental variables (temperature, pH, and $pCO_2$) across treatments. P-values represent post-hoc Tukey comparisons of generalized linear models created for a variable's effect on treatment. Statistically similar measures are highlighted in bold and indicate that the respective variable is not different between the incubators housing the contrasting treatments.*

| Treatment Comparison | Temperature p-value | pH p-value | $pCO_2$ p-value |
|---|---|---|---|
| AM – OA | **0.173416** | <0.0001 | <0.0001 |
| AM – OW | <0.0001 | 0.039984 | **0.999949** |
| AM – OWA | <0.0001 | <0.0001 | <0.0001 |
| OA – OW | <0.0001 | <0.0001 | <0.0001 |
| OA – OWA | <0.0001 | <0.0001 | **0.398534** |
| OW – OWA | **0.932459** | <0.0001 | <0.0001 |
